# A Mechanistic Modeling Framework Reveals the Key Principles Underlying Tumor Metabolism

**DOI:** 10.1101/2021.01.04.424598

**Authors:** Shubham Tripathi, Jun Hyoung Park, Shivanand Pudakalakatti, Pratip K. Bhattacharya, Benny Abraham Kaipparettu, Herbert Levine

**Affiliations:** PhD Program in Systems, Synthetic, and Physical Biology, Rice University, Houston, Texas, USA; Center for Theoretical Biological Physics and Department of Physics, Northeastern University, Boston, Massachusetts, USA; Department of Molecular and Human Genetics, Baylor College of Medicine, Houston, Texas, USA; Department of Cancer Systems Imaging, The University of Texas MD Anderson Cancer Center, Houston, Texas, USA; Dan L. Duncan Cancer Center, Baylor College of Medicine, Houston, Texas, USA

## Abstract

While aerobic glycolysis, or the Warburg effect, has for a long time been considered a hallmark of tumor metabolism, recent studies have revealed a far more complex picture. Tumor cells exhibit widespread metabolic heterogeneity, not only in their presentation of the Warburg effect but also in the nutrients and the metabolic pathways they are dependent on. Moreover, tumor cells can switch between different metabolic phenotypes in response to environmental cues and therapeutic interventions. A framework to analyze the observed metabolic heterogeneity and plasticity is, however, lacking. Using a mechanistic model that includes the key metabolic pathways active in tumor cells, we show that the inhibition of phosphofructokinase by excess ATP in the cytoplasm can drive a preference for aerobic glycolysis in fast-proliferating tumor cells. The differing rates of ATP utilization by tumor cells can therefore drive heterogeneity with respect to the presentation of the Warburg effect. Building upon this idea, we couple the metabolic phenotype of tumor cells to their migratory phenotype, and show that our model predictions are in agreement with previous experiments. Next, we report that the reliance of proliferating cells on different anaplerotic pathways depends on the relative availability of glucose and glutamine, and can further drive metabolic heterogeneity. Finally, using treatment of melanoma cells with a BRAF inhibitor as an example, we show that our model can be used to predict the metabolic and gene expression changes in cancer cells in response to drug treatment. By making predictions that are far more generalizable and interpretable as compared to previous tumor metabolism modeling approaches, our framework identifies key principles that govern tumor cell metabolism, and the reported heterogeneity and plasticity. These principles could be key to targeting the metabolic vulnerabilities of cancer.

**Author Summary:** Tumor cells exhibit heterogeneity and plasticity in their metabolic behavior, relying on distinct nutrients and metabolic pathways, and switching to reliance on different pathways when challenged by an environmental change or a drug. While multiple previous studies have focused on identifying metabolic signatures that can distinguish tumor cells from non-tumorigenic ones, frameworks to analyze the metabolic heterogeneity in tumors have been lacking. Here, we present a mechanistic mathematical model of some of the key metabolic pathways active in tumor cells and analyze the steady state behaviors the model can exhibit. We find that the rate of ATP use by tumor cells can be a key determinant of the metabolic pathway via which tumor cells utilize glucose. We further show that tumor cells can utilize different pathways for satisfying the same metabolic requirements, and explore the implications of such behavior for the response of tumor cells to drugs targeting tumor metabolism. At each step, we discuss how our model predictions fit within the context of experimental observations made across tumor types. The present modeling framework represents an important step towards reconciling the wide array of experimental observations concerning tumor metabolism, and towards a more methodical approach to targeting tumors’ metabolic vulnerabilities.

## Introduction

Proposed as an emerging hallmark of cancer nearly a decade ago [1], metabolic reprogramming has now entered into the limelight of cancer biology as a key feature of tumor cells across cancer subtypes, with multiple therapeutic implications [2]. Aerobic glycolysis, commonly known as the Warburg effect [3], characterized by increased glucose uptake most of which is excreted out as lactate even under normoxic conditions, has been synonymous with cancer cell metabolism for nearly a century. Studies carried out over the past decade have however revealed a more complex picture— there exists widespread intra-tumoral heterogeneity not only in the way tumor cells utilize glucose but also in the activities of the various other metabolic pathways in tumor cells. Further, tumor cells at different stages of metastatic progression exhibit different metabolic phenotypes and metastases growing in distinct organs can also exhibit differences in their metabolic activity [4,5]. Understanding the mechanistic basis of metabolic heterogeneity in different contexts will be key to the design of anti-cancer metabolic therapies.

**Figure 1.**
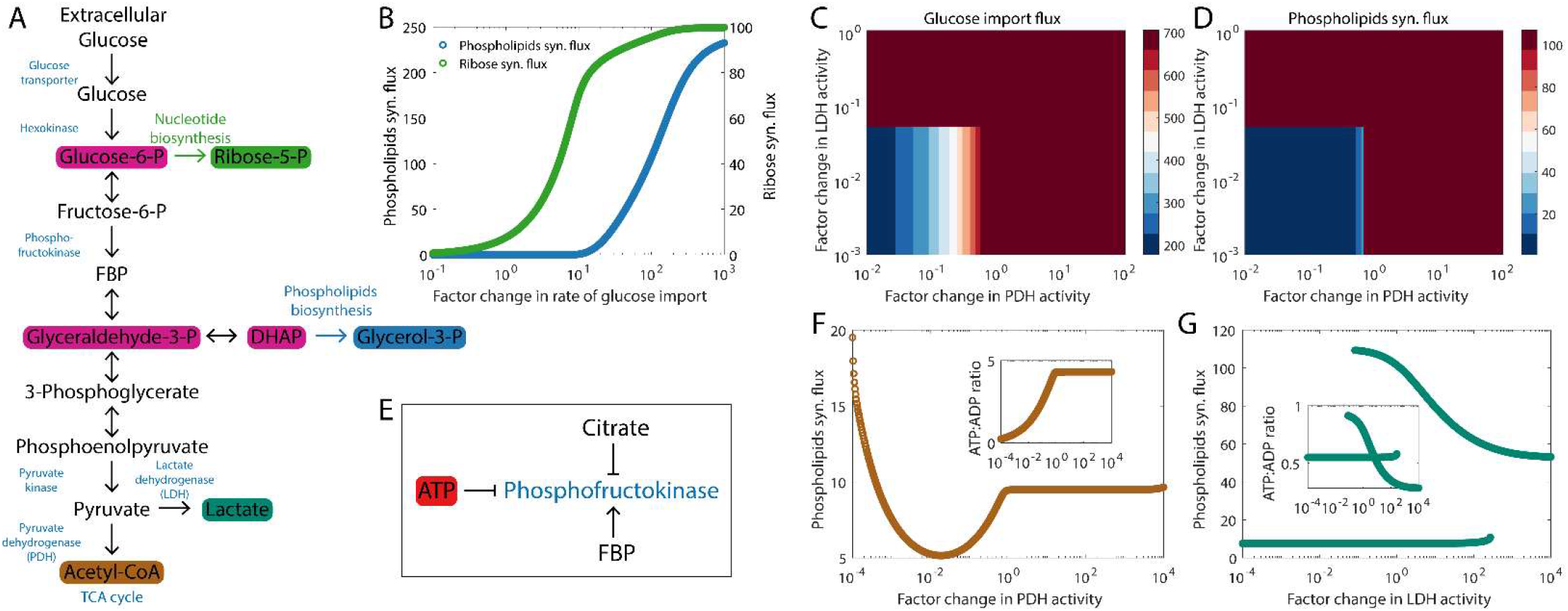
High ATP production during oxidative phosphorylation can drive a preference for aerobic glycolysis in fast proliferating cells. (A) Reactions in the glycolytic pathway and key anabolic processes that use glycolytic intermediates as substrates. The pyruvate generated at the end of glycolysis can either be excreted out as lactate or enter the TCA cycle as acetyl-CoA. (B) Increased glucose uptake can drive large fluxes through the ribose synthesis pathway and through the phospholipids synthesis pathway. (C) Increasing the activity of pyruvate dehydrogenase (PDH), lactate dehydrogenase (LDH), or both can increase the glucose uptake. (D) The phospholipids synthesis flux increases upon the upregulation of PDH, LDH, or both. In (C) and (D), we have assumed that the ATP concentration in the cytoplasm remains constant despite the higher ATP production per glucose molecule when glucose is allowed to enter the TCA cycle by upregulating PDH activity. (E) Different metabolites that can modulate the enzymatic activity of phosphofructokinase (PFK). FBP: fructose-1, 6-biphosphate. (F) Variation of the phospholipids synthesis flux with an increase in PDH activity (while keeping LDH activity fixed). (G) Variation of the phospholipids synthesis flux with an increase in LDH activity (while keeping PDH activity fixed). The insets in (F) and (G) show how the ATP:ADP ratio varies in each case. The bistability in the phospholipids synthesis flux as the LDH activity is varied arises from a positive feedback loop— fructose-1, 6-biphosphate, the product of PFK’s enzymatic activity, can allosterically activate the enzyme (see panel E). Fluxes are in units of mM h^-1^ (millimolar per hour) and are shown at steady state.

A mechanistic modeling framework to understand the mechanistic underpinnings of tumor metabolic heterogeneity has been lacking. Multiple previous studies have relied upon constraint-based models such as flux balance analyses, often involving genome-scale metabolic models, to simulate the metabolic response of tumor cells and to identify the metabolic vulnerabilities (reviewed in [6]). Among the studies focused on explaining the Warburg effect, Vazquez *et al*. [7] showed that a switch to aerobic glycolysis can maximize the rate of ATP production by cancer cells under high glucose uptake while Shlomi *et al*. [8] have shown that under a solvent capacity constraint, the Warburg effect maximizes the rate of biomass production in cancer cells. Constraint-based modeling has been a popular approach for identifying anti-cancer targets [9–11] and for proposing cancer-specific metabolic signatures [12–15]. The analyses in these studies do not predict the widespread heterogeneity in the metabolic profiles of cancer cells in the primary tumor and of tumor cells at different stages of metastatic disease progression, mainly due to the involvement of a global objective function that all tumor cells must optimize. Further, the frameworks in these studies do not connect the metabolic heterogeneity in tumor cells to the heterogeneity in other phenotypic states such as cell migration. While mechanistic models of tumor metabolism are more suited to addressing the questions concerning metabolic heterogeneity, past analysis has instead focused on fitting detailed models to a specific experimental setup [16] instead of identifying the general principles underlying tumor metabolism.

Recently, Jia *et al*. have put forth a systems-level analysis of coupled gene regulatory networks and metabolic pathways to describe metabolic heterogeneity and plasticity in tumor cells [17]. The study shows that cancer cells can switch between different metabolic states in response to changes in the activities of master regulators such as AMPK and HIF-1. However, the modeling approach therein is too coarse-grained to answer some interesting questions relating to the flux through different reactions in the metabolic pathways and to predict the detailed cellular response to perturbations in the activities of specific enzymes.

Here, we construct a mechanistic model which incorporates the key metabolic reactions that have been shown to be active in cancer cells. Instead of relying on a stoichiometric modeling framework such as flux balance analysis [18], we write down detailed mathematical equations describing the kinetics of different enzymatic reactions. The resultant system of ordinary differential equations can be numerically integrated to determine the steady state of the metabolic system. We show that the preference for aerobic glycolysis increases the flux through the anabolic pathways that use glycolytic intermediates as substrates, thereby facilitating fast proliferation. This results from the low ATP production when glucose is excreted out as lactate as compared to when glucose enters the TCA cycle as pyruvate. The rate of ATP consumption in cancer cells, which can vary with other cellular phenotypic properties, can thus modulate the preference of cancer cells for aerobic glycolysis versus oxidative phosphorylation. Next, we explore how the relative dependence of cancer cells on glucose and glutamine and the relative availability of these nutrients in the microenvironment can affect the metabolic profiles of tumor cells. We further use the model to predict the changes in the metabolic and gene expression profiles of melanoma cells under treatment with a BRAF inhibitor that can suppress glutamine uptake. Finally, we discuss how different experimental observations relating to cancer cell metabolism fit within the context of our modeling framework.

## Results

### Developing a mechanistic framework to model tumor metabolism

Our goal, in this study, is to explore the range of dynamical behaviors that can be exhibited by a new mathematical model which incorporates the critical aspects of tumor metabolism. We then analyze if the observed model behaviors can qualitatively explain any of the reported metabolic behaviors of tumor cells. This approach is in contrast to previous mechanistic modeling approaches that focus on fine tuning model parameters so that the model behavior is in agreement with the observations from a specific experimental setup (see, for example, Roy and Finley [16]), and allows us to develop a framework that can be utilized to analyze the metabolic behavior of tumor cells across cancer types and in different environments. The present framework is motivated by the success of a similar strategy in modeling the phenotypic heterogeneity and plasticity arising from gene regulatory networks [19,20].

Our model of tumor metabolism was constructed using the information available from the literature regarding the enzymatic reactions involved in key metabolic pathways active in tumor cells. These include glycolysis, the TCA cycle, oxidative phosphorylation, and glutaminolysis. A similar approach— obtaining the subset of cancer-specific metabolic reactions from the generic set of human metabolic reactions— has been popular for the construction of genome-scale models of tumor metabolism [9,12,13]. Here, instead of choosing the cancer-specific metabolic reactions on the basis of cancer cell proteomics [9,12] or gene expression data [13], we chose the metabolic pathways for analysis on the basis of the available literature on tumor metabolism (reviewed in [21]). The dynamics of the concentrations of different metabolites in our setup were modeled using ordinary differential equations. The mathematical expressions describing the activities of the different enzymes and the relevant kinetic parameters were taken from the literature [22–26]. Oxidative phosphorylation and the electron transport chain reactions were treated using a simplified model proposed previously by Nazaret *et al*. [27]. Activation and inhibition of different enzymes were modeled by changing the reaction velocity (*V_max_*) for the enzyme by a multiplicative factor. The differential equations in the modeling setup (included in the S1 Text) were integrated numerically to determine the steady state model behavior.

A key challenge in the mechanistic modeling of cell metabolism is the unavailability of many of the mathematical expressions describing the kinetic behavior of different enzymes and the relevant kinetic parameters. In such cases, we made reasonable assumptions based on Michaelis-Menten kinetics [28]. Even in cases where the relevant kinetic information is available, the data obtained from different sources are often not compatible [23]— the expressions describing enzyme kinetics as well as the relevant kinetic parameters used in the present study were obtained from multiple previous studies [22–26] which themselves relied on results from experiments conducted in different tissue types, in cells from different species, and under a varied set of conditions. The incompatibility of the kinetic parameters can lead to scenarios wherein the different metabolic fluxes do not balance, and the system does not exhibit a non-trivial bounded steady state. To address this issue, some of the kinetic parameters obtained from previous studies were manually adjusted until the model exhibited a bounded steady state. This constraint on the model kinetic parameters is similar to the constraint on the steady state metabolic fluxes commonly imposed during flux balance analyses [18]. The values of the different kinetic parameters are included in the S1 Text. The S1 Text also includes a more detailed discussion of the assumptions underlying model construction. Matlab code used to simulate the model behavior is available online (https://github.com/st35/cancer-metabolism).Note that the present modeling framework is not being put forth as an alternative to genome-scale metabolic models which allow for a global analysis of the role of different enzymes and metabolic pathways in tumor metabolism. Instead, the focus of the present model is on understanding the more microscopic picture— the role of metabolic feedback loops and of different enzymes in specific contexts. The model sacrifices completeness for interpretability and mechanistic insight.

In the following subsections, we discuss the behaviors exhibited by our metabolic model and their relevance to cancer biology.

### Inhibition of phosphofructokinase by ATP can drive the preference for aerobic glycolysis in fast proliferating cells

As shown in Fig. 1 A, intermediate metabolites generated during the multi-step process that converts glucose to pyruvate are used in key anabolic processes required for cell division [21]. These include the ribose synthesis pathway, crucial for nucleotide synthesis, and the phospholipids synthesis pathway. A large flux through both these pathways is likely to facilitate fast proliferation. To determine how this may be achieved, we simulated the dynamics of the glycolysis pathway along with the first reactions in the ribose synthesis and the phospholipids synthesis pathways. Fig. 1 B shows that the steady state flux through both ribose synthesis and phospholipids synthesis pathways can be increased by increasing the rate of glucose uptake. To obtain this result, we have assumed that all the pyruvate that is generated from the glucose taken up by a cell is utilized via some metabolic process inside the cell and that any ATP produced during such a process is also used up, thereby keeping constant the concentration of ATP inside the cell. The pyruvate generated from glucose can have two primary fates (Fig. 1 A). On the one hand, lactate dehydrogenase (LDH) can convert pyruvate into lactate which is then excreted out of the cell. On the other hand, the pyruvate can enter the mitochondria where it is converted to acetyl coenzyme A (acetyl-CoA) by the enzyme pyruvate dehydrogenase (PDH) and further oxidized via the TCA cycle. The rate of glucose uptake in tumor cells can be increased by increasing the activity of either of the two enzymes (Fig. 1 C). The increased glucose uptake in turn increases the flux through the phospholipids synthesis pathway (Fig. 1 D). The symmetry between increasing glucose uptake and the phospholipids synthesis flux via increased lactate excretion and via increased flux through the TCA cycle, apparent in Fig. 1 C and 1 D, is broken when the rates of ATP production in the two processes are taken into consideration (Fig. 1 E-G). If the carbon taken in as glucose is excreted out as lactate, 2 ATP molecules per molecule of glucose are produced. Using the glucose carbon to drive the TCA cycle followed by oxidative phosphorylation can generate 30-32 molecules of ATP per glucose molecule [29]. At a fixed rate of ATP consumption by the cell, using a large fraction of the glucose taken up by the cell to drive the TCA cycle can lead to increased ATP accumulation in the cytoplasm (see inset in Fig. 1 F) which will inhibit the glycolytic enzyme phosphofructokinase (PFK) (Fig. 1 E), shutting down glucose uptake [30] and driving down the phospholipids synthesis flux (Fig. 1 F). Excreting out the glucose carbon as lactate, on the other hand, can limit the ATP-mediated downregulation of PFK activity (see inset in Fig. 1 G), thereby maintaining a high phospholipids synthesis flux (Fig. 1 G) which would help maintain fast proliferation rates. This offers a possible explanation for the preference for glucose carbon secretion as lactate even under normoxic conditions (the Warburg effect) seen in fast proliferating tumor cells across cancer types.

### Varying ATP requirements can drive heterogeneity in the metabolic phenotype exhibited by tumor cells

In the previous section, we have shown that the inhibition of PFK due to the accumulation of excess ATP (generated by allowing glucose-derived pyruvate to enter the TCA cycle) can downregulate anabolic processes including phospholipids synthesis. We propose that this could be the reason many fast proliferating cells exhibit the Warburg effect. The hypothesis implies that modulation of the rate of ATP consumption in cells can change which metabolic phenotype will lead to a large flux through anabolic processes and thus facilitate fast proliferation. Fig. 2 A shows that the metabolic phenotype that maximizes the flux through the phospholipids synthesis pathway can change as the rate of ATP consumption by tumor cells changes. At low, basal rates of ATP consumption, the high lactate secretion, low TCA cycle flux (high LDH, low PDH) metabolic phenotype is needed for driving a large flux through the phospholipids synthesis pathway. A switch to a phenotype with high TCA cycle flux (high PDH) will shut down this key anabolic process. However, in cells that consume ATP at very high rates, for example, in metastasizing cancer cells that are actively migrating through the extracellular matrix [31], ATP will not accumulate to concentrations high enough to inhibit PFK activity even when large amounts of ATP are produced per glucose molecule via the TCA cycle. These cells can thus sustain fast proliferation rates while not exhibiting the Warburg effect. Difference in ATP consumption by cells at different stages of metastasis can thus contribute towards the differences in the metabolic profiles of tumor cells at distinct stages of metastasis. Note that in Fig. 2 A, we have assumed that there is sufficient oxygen available for ATP generation via the TCA cycle followed by oxidative phosphorylation. When the oxygen supply is limited (as is often the case in the interior of solid tumors), cells have to rely solely on converting glucose to lactate for ATP production (anaerobic glycolysis). In such a scenario, the rate of lactate production will be higher in cells with higher rates of ATP consumption (Fig. 2 B).

**Figure 2.**
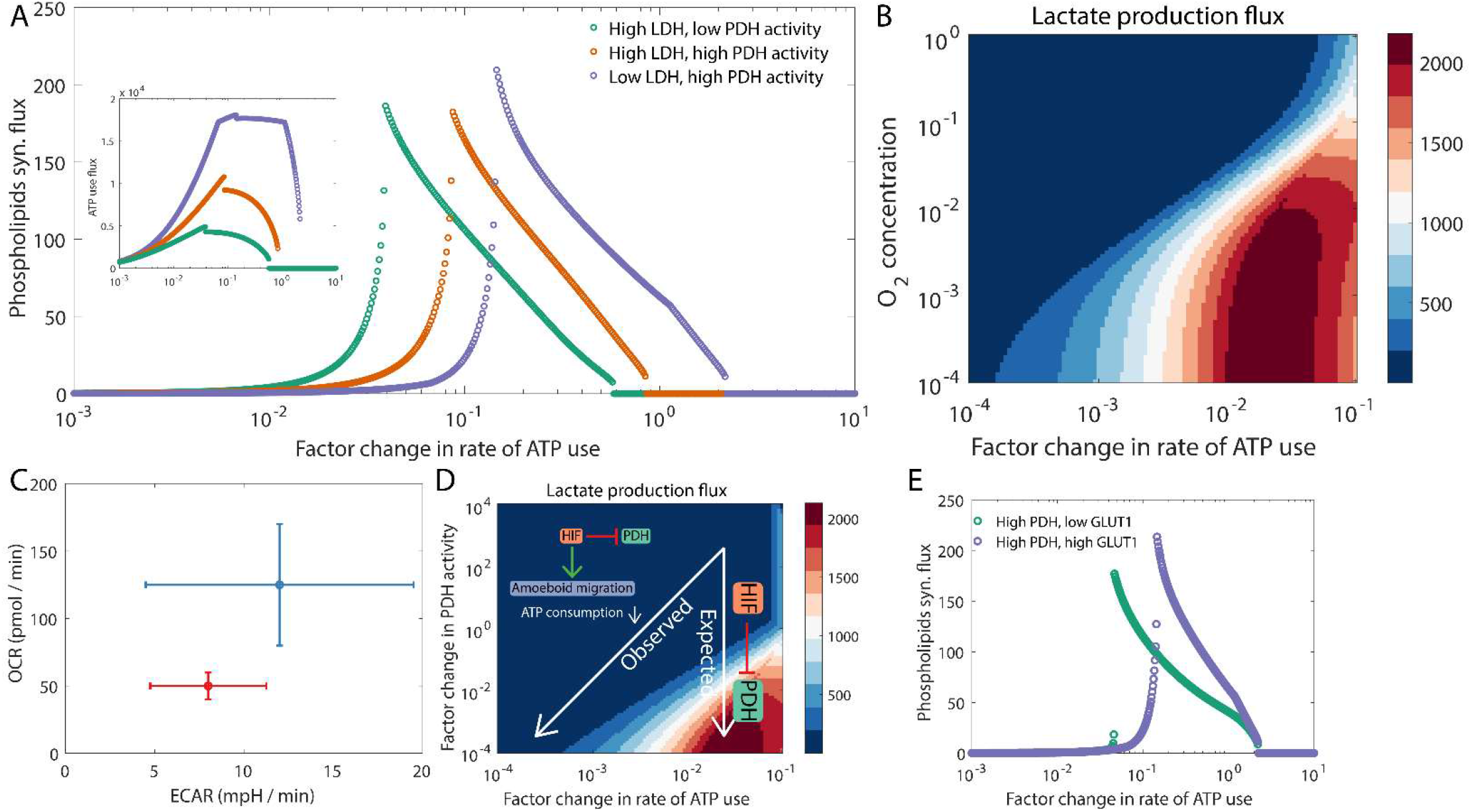
Rate of ATP consumption by tumor cells can modulate the preference for aerobic glycolysis versus oxidative phosphorylation. (A) The regime that maximizes the phospholipids synthesis flux depends on the rate of ATP consumption. The inset shows that the flux through pathways that consume ATP is maximized when the activity of PDH is high, i.e., when most of the glucose carbon enters the TCA cycle. (B) The rate of lactate production by tumor cells can depend not only on the availability of oxygen in the microenvironment but also the rate of ATP consumption. (C) Change in the metabolic profile of 4T1 cells upon HIF1 stabilization (red) as compared to the control (blue). The behavior shown is representative of that reported by te Boekhorst *et al*. [32] (see Fig. 5e therein). ECAR: extracellular acidification rate, which correlates positively with the rate of lactate excretion by cells. (D) Change in the metabolic profile of cells upon HIF1 stabilization (which inhibits PDH activity) as predicted by our model. The change labeled as expected is when the migratory phenotype (and the rate of ATP consumption) remains unchanged upon HIF1 stabilization. The change labeled as observed is when HIF1 stabilization is accompanied by a switch from actin-driven migration to amoeboid migration [32] which decreases the rate of ATP consumption during cell migration. (E) Under high PDH activity and very high ATP consumption rate, GLUT1 up-regulation can increase the phospholipids synthesis flux. Fluxes are in units of mM h^-1^ (millimolar per hour) and are shown at steady state.

A connection between ATP consumption by tumor cells and their metabolic phenotype was recently demonstrated by te Boekhorst *et al*. [32]. They observed that the stabilization of HIF1 (which downregulates PDH activity) in 4T1 mouse breast cancer cells decreased the oxygen consumption rate (OCR), as expected, but surprisingly did not significantly increase the rate of lactate production (Fig. 2 C). The decrease in OCR is indicative of a decrease in the flux through the TCA cycle. Our model predicts that this can happen if ATP consumption by tumor cells also decreases along with the downregulation of PDH activity upon HIF1 stabilization (Fig. 2 D). This is indeed true in the case of 4T1 cells— upon DMOG treatment (which stabilizes HIF1), 4T1 cells underwent a change in their mode of migration from an actin-driven mode to an amoeboid mode involving blebbing protrusions [32]. ATP consumption in the actin-driven mode of cell migration is much higher as compared to that in the cell migration mode dominated by blebbing protrusions [31]. The decrease in ATP consumption upon the switch in the migration mode could have suppressed the expected increase in lactate production in this experiment (Fig. 2 D). Whether the switch in the mode of migration is driven by changes in gene expression caused directly by HIF1 stabilization or by the metabolic changes in response to HIF1 stabilization remains a very interesting but yet unanswered question.

Another scenario wherein differing ATP requirements can underlie the observed metabolic heterogeneity is collective invasion by tumor cells. Leader cells, which are at the invading edge of the tumor cell pack and use ATP for membrane protrusion, for maintaining focal adhesion stability, and for remodeling the extracellular matrix, have a higher ATP requirement as compared to the trailing follower cells. To maximize ATP production, leader cells must rely on the TCA cycle and oxidative phosphorylation which help drive a large flux through any pathway that consumes ATP (see inset in Fig. 2 A). Our model predicts that given the reliance of leader cells on the TCA cycle, they will likely proliferate at a rate lower than that of follower cells (which exhibit high lactate production) for a range of ATP consumption rates (between 10^-2^ and 10^-1^ on the horizontal axis in Fig. 2 A, for example). Such behavior was recently observed by Commander *et al*. [33] for the H1299 non-small cell lung cancer cell line— follower cells isolated from the cultures of this cell line exhibited higher lactate production and higher proliferation rates as compared to the leader cells from the same culture. Our model further predicts that under very high rates of ATP consumption when complete glucose oxidation via the TCA cycle (driven by high PDH1) is the preferred metabolic state, the phospholipids synthesis flux will increase upon the upregulation of GLUT1 (a glucose transporter) activity (Fig. 2 D). The observation that GLUT1 overexpression in leader cells can increase their proliferation rate [33] provides some preliminary evidence in support of this prediction.

The results reported in this section illustrate how our modeling framework can be used to explain the co-variation of such seemingly disparate phenotypes as cell proliferation and cell migration. Note that here we have focused on the phospholipids synthesis pathway as a representative example from the set of anabolic pathways that use glycolytic intermediates as substrates. The qualitative model predictions will remain unchanged for any other anabolic pathway of interest which relies upon glycolytic intermediates generated downstream from the enzyme PFK (for example, the serine synthesis pathway [34]).

### Distinct anaplerotic pathways can contribute towards *de novo* fatty acid synthesis

Fatty acids are a key biomolecular requirement for membrane synthesis, a necessity for cell division. While some tumor cells can take up fatty acids from the extracellular environment, others must synthesize the required fatty acids ***de novo*** [35]. A key substrate for fatty acid synthesis is acetyl-CoA, an intermediate of the TCA cycle, which is removed from the TCA cycle as citrate (Fig. 3 A). In the cytoplasm, the enzyme ACLY converts citrate back to acetyl-CoA. The enzyme ACC then converts acetyl-CoA to malonyl-CoA, a precursor for the synthesis of fatty acids [35]. To keep the sequence of reactions in the TCA cycle going, the intermediates must be replenished to compensate for the loss of citrate. The metabolic reactions that replenish the metabolites harvested from the TCA cycle for other cellular processes are called anaplerotic reactions [36]. Glutamine is a key biomolecule that can contribute towards anaplerosis during ***de novo*** fatty acid synthesis— glutamine can be converted to α-ketoglutarate which replenishes the carbon removed from the TCA cycle as citrate (Fig. 3 A). As shown in Fig. 3 B, when the fatty acid synthesis pathway is active (high ACC activity), our model predicts that the TCA cycle will shut down in the absence of glutamine unless there are other active pathways that can replenish the TCA cycle carbon lost as citrate. Sufficient glutamine availability is needed to drive a large flux through the fatty acid synthesis pathway (Fig. 3 C). Additionally, oxygen is essential for the oxidation of NADH via oxidative phosphorylation and the electron transport chain to keep the TCA cycle running. Therefore, the anaplerotic pathway described above can only function when oxygen is available (green curve in Fig. 3 E) since it involves all the reactions of the TCA cycle. Under hypoxic conditions, an alternate reaction pathway, mediated by NADPH-dependent IDH, may be activated to utilize glutamine for anaplerosis— reductive carboxylation of α-ketoglutarate to directly form citrate (Fig. 3 D and orange curve in Fig. 3 E). This model behavior is in agreement with the experimental observation that hypoxia can reprogram A549 lung cancer cells to rely on reductive glutamine metabolism, and inhibition of NADPH-dependent IDH can suppress the proliferation of these cells [37]. Overall, our model shows that depending on the oxygen availability in different parts of the tumor microenvironment, proliferating cells can exhibit varying activity levels of different glutamine-driven anaplerotic pathways. Metabolic profiles of cells proliferating under normoxic and hypoxic conditions in a glutamine-dependent manner will therefore be different (Fig. 3 F).

**Figure 3.**
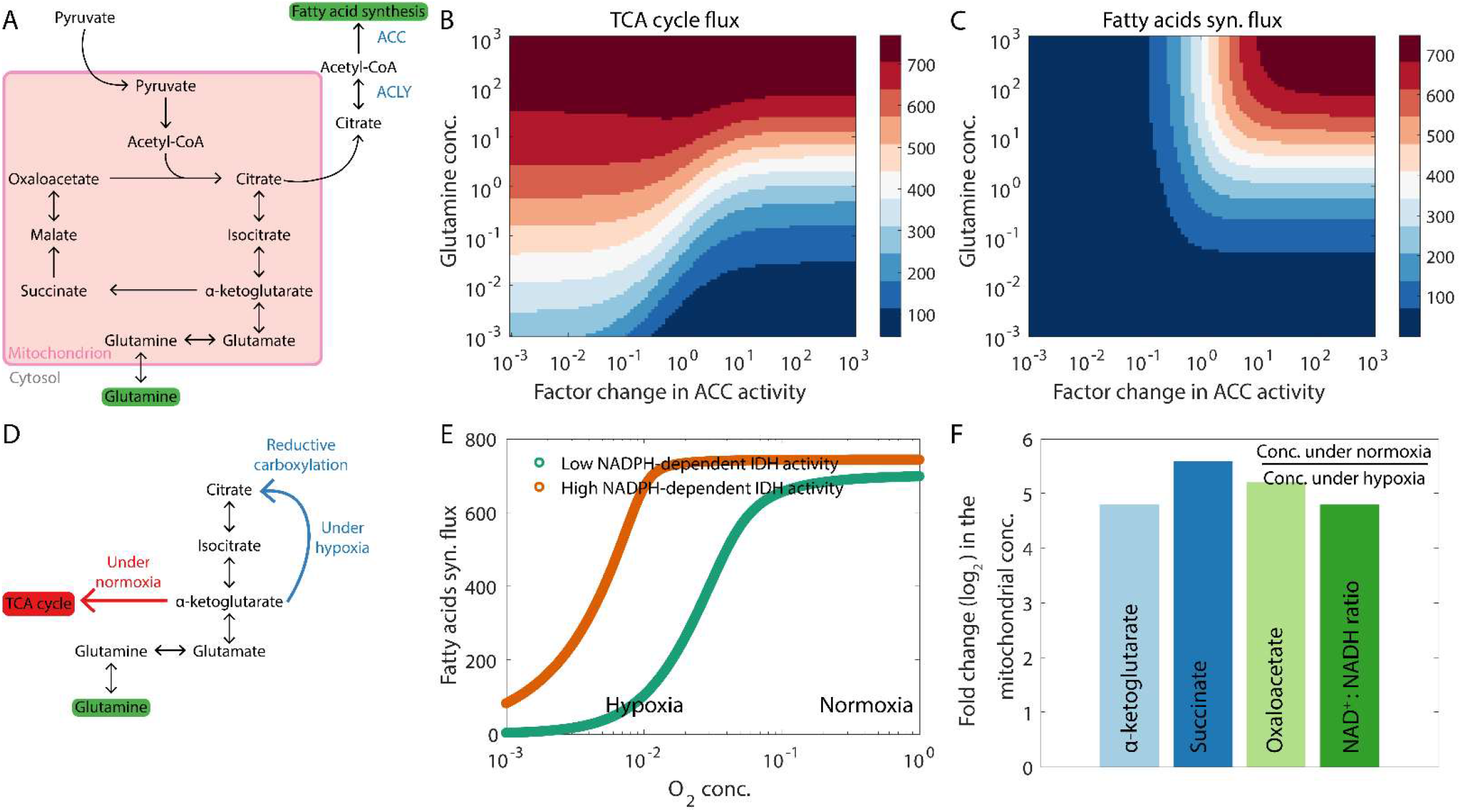
Glutamine is a key metabolite for anaplerosis during de novo fatty acid synthesis and can be utilized via different pathways under different conditions. (A) Reactions in the TCA cycle. The citrate removed from the TCA cycle for use as a substrate for fatty acid synthesis is replenished by the α-ketoglutarate generated from glutamine. (B) At high rates of de novo fatty acid synthesis (indicated by high ACC activity), our model predicts that the TCA cycle will shut down at low glutamine concentrations. Here, the flux through the enzyme α-ketoglutarate dehydrogenase is shown as the TCA cycle flux. (C) High glutamine availability can drive a large fatty acid synthesis flux. (D) Glutamine can be utilized for fatty acid synthesis via distinct pathways under normoxic and hypoxic conditions. (E) Under hypoxic conditions, glutamine can be utilized via the reductive carboxylation pathway (see panel D). High activity of NADPH-dependent IDH is essential for reductive carboxylation. (F) Ratio of the concentration of different TCA cycle intermediates under normoxic conditions to the concentration under hypoxic conditions. Also shown is the difference in the mitochondrial NAD^+^:NADH ratio under the two conditions. Since the NADPH-dependent IDH pathway (active under hypoxic conditions) skips the TCA cycle reactions, activation of this pathway is accompanied by a decrease in the mitochondrial concentrations of multiple TCA cycle intermediates. Fluxes are in units of mM h^-1^ (millimolar per hour) and are shown at steady state.

Under glutamine deprivation due to a lack of glutamine availability in the microenvironment or due to treatment with a drug that inhibits glutamine uptake, a glutamine-independent anaplerotic pathway will be needed for ***de novo*** fatty acid synthesis. Pyruvate carboxylase (PC) can drive one such pathway which involves the conversion of pyruvate to oxaloacetate (Fig. 4 A). As shown in Fig. 4 B, under conditions of glutamine deprivation, cells with high PC activity can drive fatty acid synthesis by generating both the precursors of citrate— oxaloacetate and acetyl-CoA— from glucose-derived pyruvate. Our model predicts that cells with low PC activity cannot synthesize fatty acids ***de novo*** under glutamine deprivation and are thus more likely to be sensitive to glutamine deprivation therapies. Consistent with our model prediction, MC-38 mouse colon cancer cells, unable to upregulate PC activity, have been shown to be susceptible to glutamine blockade. We note in passing that in the same experimental setup, T cells showed increased PC activity upon glutamine deprivation and maintained a proliferative phenotype under glutamine blockade by using glucose for anaplerosis [38]. Note that the PC-driven anaplerotic process short-circuits the TCA cycle (Fig. 4 A). Cells relying on this pathway will thus likely exhibit low TCA cycle flux and, consequently, a low oxygen consumption rate (Fig. 4 C) as compared to cells that rely on oxidative glutamine metabolism. Moreover, these cells will also exhibit low rates of lactate production as compared to glutamine-dependent cells since double the number of pyruvate molecules is now needed for citrate synthesis (Fig. 4 D), decreasing the availability of pyruvate for lactate production.

**Figure 4.**
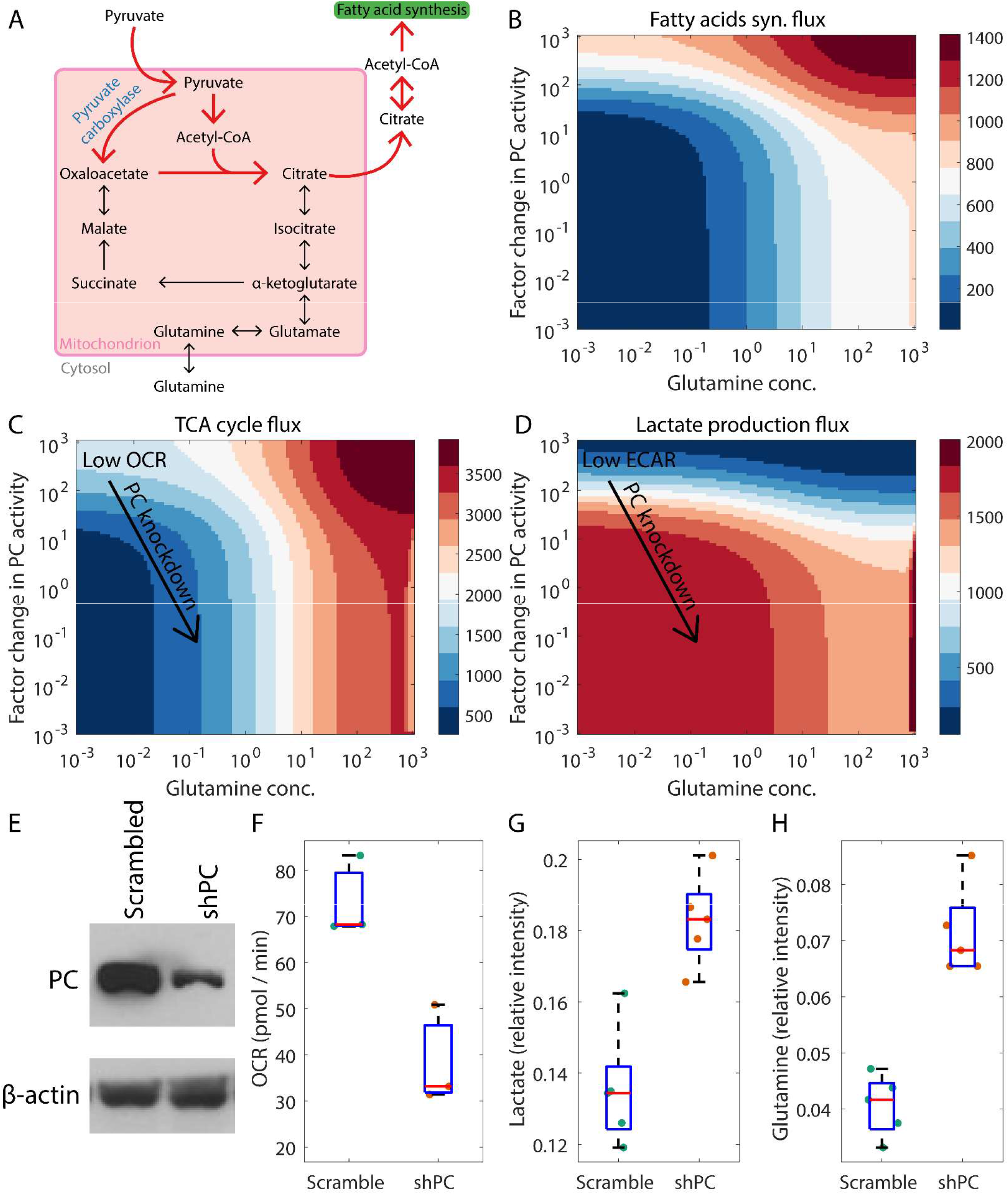
Pyruvate carboxylase (PC)-dependent anaplerosis under low glutamine concentrations. (A) Reactions involved in PC-dependent anaplerosis during fatty acid synthesis. (B) At low glutamine concentrations, high PC activity can help maintain a significant fatty acid synthesis flux. (C) and (D) When PC-dependent anaplerosis is the dominant anaplerotic pathway, cells carrying out ***de novo*** fatty acid synthesis will exhibit low TCA cycle flux and low oxidative phosphorylation (and consequently, low OCR) (C) and low lactate production (and consequently, low ECAR) (D). ECAR: extracellular acidification rate, which correlates positively with the rate of lactate excretion by cells. In panel C, the rate of conversion of TCA cycle-generated NADH to NAD^+^ in an oxygen-dependent manner is shown as the TCA cycle flux. (E-G) Downregulating PC expression in MDA-MB-468 breast cancer cells decreases the oxygen consumption rate (OCR) and upregulates lactate production. (E) Western blot confirmed a decrease in the expression of the PC protein upon knockdown using PC shRNA compared to cells with control (Scrambled) shRNA. (F) Seahorse analysis shows decreased OCR in shPC cells (also, see arrow in panel C). NMR spectroscopy analysis shows increased lactate level (G) (also, see arrow in panel D) and increased glutamine level (H) in shPC cells. Fluxes are in units of mM l^-1^; (millimolar per hour) and are shown at steady state (B-D).

To determine if PC activity is indeed associated with low lactate production as per the above prediction, we used a short hairpin RNA (shPC) to downregulate the expression of PC in MDA-MB-468 breast cancer cells (Fig. 4 E). Arrows in Fig. 4 C and Fig. 4 D indicate that downregulation of PC activity should be accompanied by a decline in the TCA cycle flux (and consequently, a decrease in the basal oxygen consumption rate (OCR)) and an increase in lactate production. This was confirmed by Seahorse analysis (Fig. 4 F) and NMR spectrometry (Fig. 4 G), respectively. Interestingly, PC knockdown was also accompanied by an increase in cellular glutamine concentration (Fig. 4 H), indicating that increased glutamine uptake may compensate for decreased PC activity in MDA-MB-468 cells. Whether the increase in glutamine uptake upon PC knockdown is driven purely by metabolic feedback or by some gene regulatory mechanism remains to be determined.

Finally, we examined the model behavior under glucose deprivation and sufficient glutamine availability. As shown in Fig. 1 A and Fig. 5 C, cells are unlikely to be able to proliferate under such conditions [39]. This is because key anabolic processes that use glycolytic intermediates as substrates, such as the ribose synthesis and the phospholipids synthesis pathways, will shut down when glucose is unavailable. However, cells can survive in such a scenario provided sufficient ATP is available to keep the basal cellular processes going. ATP can be generated in the absence of glucose by utilizing glutamine to drive the TCA cycle (Fig. 5 A). The enzyme ME2 (malic enzyme 2), which converts malate directly to pyruvate, can help drive a glutamine-dependent TCA cycle. Cells with high ME2 activity can thus survive glucose deprivation by using glutamine to generate ATP (Fig. 5 B). In agreement with our model prediction, inhibition of both mitochondrial pyruvate import and the enzyme glutamate dehydrogenase (a key enzyme for generating α-ketoglutarate from glutamine) has been shown to be cytotoxic in glioblastoma cells as compared to only inhibiting mitochondrial pyruvate import [40]. Note that ATP synthesis from glutamine in a glucose-independent manner is oxygen-dependent (Fig. 5 D)— the process involves an operational TCA cycle which generates NADH, and the conversion of glutamate to α-ketoglutarate generates an additional NADH molecule. Oxygen must be available for the oxidation of NADH to NAD^+^ via the electron transport chain, a process that generates ATP. Under hypoxic conditions, NADH will accumulate and shut down not only the TCA cycle but also the entry of glutamine into the TCA cycle (Fig. 5 D). Cells in hypoxic niches are thus less likely to survive glucose deprivation as compared to cells in normoxic niches.

**Figure 5.**
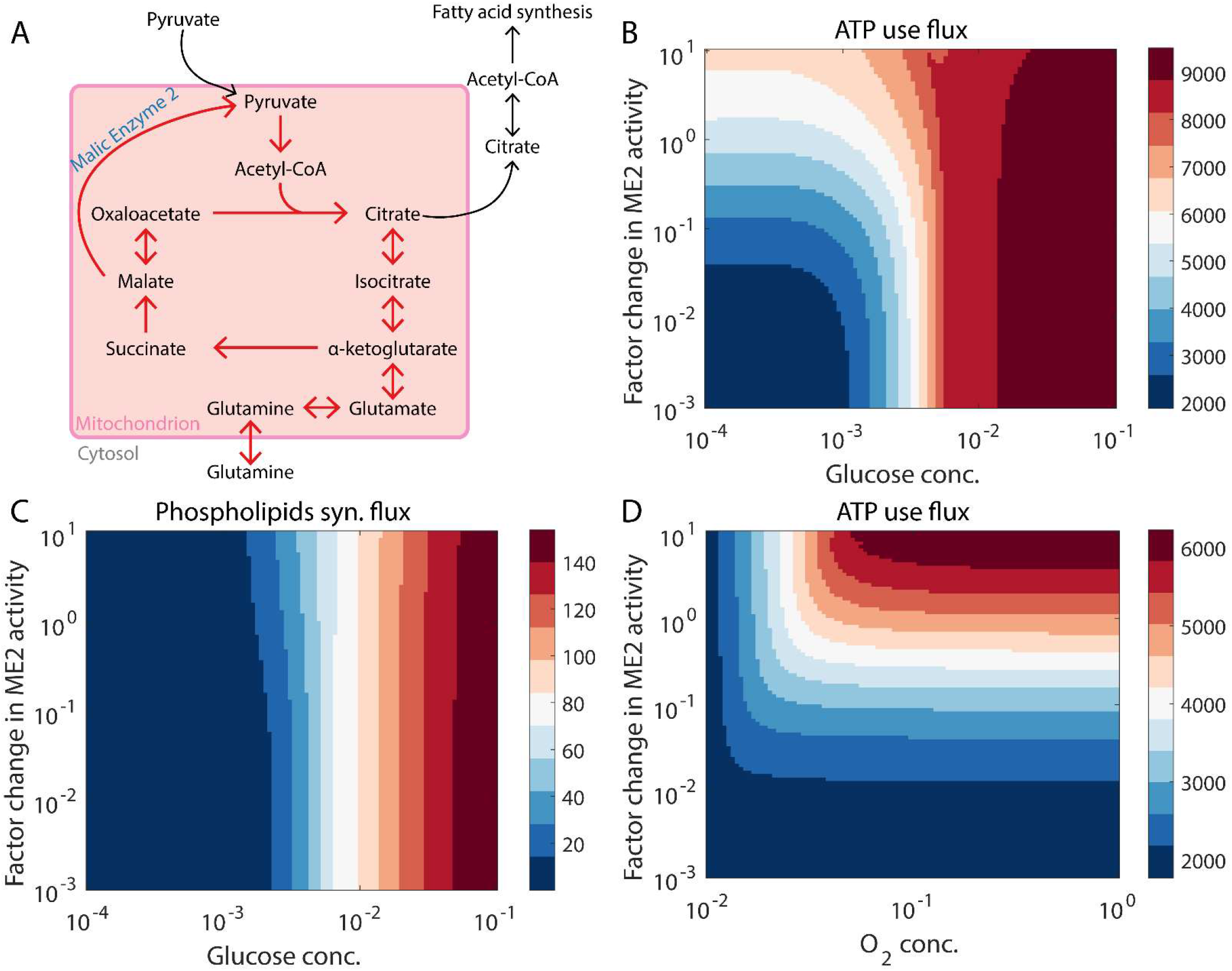
Glutamine can drive the TCA cycle in the absence of glucose. (A) Reactions involved in the glutamine-driven TCA cycle when glucose concentration is low. (B) Under low glucose concentrations, malic enzyme 2 (ME2) activity must be upregulated to maintain a significant ATP production rate. (C) In the absence of glucose, the phospholipids synthesis flux remains low, irrespective of the ME2 activity. Thus, cells are unlikely to proliferate under low glucose concentrations [39]. (D) Oxygen is essential for maintaining a significant flux through pathways that consume ATP when glutamine is available, but the glucose concentration is low. Fluxes are in units of mM h^-1^ (millimolar per hour) and are shown at steady state.

### Interplay between metabolic plasticity and drug response in melanoma

Paudel *et al*. [41] observed that single-cell-derived subclones of the SKMEL5 melanoma cell line, which carries a BRAF mutation, exhibit heterogeneity in their short-term response to the BRAF inhibitor PLX4720 (Fig. 6 A; also see Fig. 2 A and Fig. 3 A in Paudel *et al*. [41]). Since BRAF signaling inhibition can suppress glutamine uptake by downregulating MYC [44,45], cells proliferating under PLX4720 treatment may resort to glutamine-independent anaplerotic pathways such as the PC-mediated pathway described above. We analyzed the gene expression profiles of the SKMEL5 subclones (reported by Jia *et al*. [42]) and observed that the expression of PC is upregulated in the SC10 subclone upon PLX4720 treatment (Fig. 6 B). Surprisingly, among the three subclones for which gene expression data is available, only SC10 showed population expansion in the short-term (< 150 hours) after PLX4720 treatment (Fig. 6 A). While subclone SC07 population is stationary, subclone SC01 exhibits population regression upon PLX4720 treatment. Both these subclones did not exhibit increased PC expression upon drug treatment (Fig. 6 B). Our modeling framework predicts that if SC10 cells are reliant on PC for anaplerosis under PLX4720 treatment as PC upregulation upon drug treatment would suggest, they must exhibit a low oxygen consumption rate (low OCR) and a low extracellular acidification rate due to low lactate production (low ECAR) (Fig. 4 C). Indeed, upon PLX4720 treatment, SC10 cells have been shown to switch to a metabolic state with low OCR and low ECAR (see Fig. 2 D in Jia *et al*. [42]). We further analyzed the gene expression profiles of melanoma tissue samples obtained from patients before and after treatment with a BRAF signaling inhibitor [43] (Gene Expression Omnibus [46] accession GSE75299). Out of the six patient samples, four exhibited upregulation of PC upon drug-induced BRAF signaling inhibition (Fig. 6 C). This is accompanied by the downregulation of MYC, a well-known target of BRAF signaling. Thus, our analysis suggests, albeit preliminarily, that PC-mediated anaplerosis may play a key role in the response of melanoma cells to treatment with BRAF signaling inhibitors, at least at short time scales. One immediate consequence of this observation is the suggestion that co-treatment with inhibitors of BRAF and PC could be a promising anti-melanoma therapy.

**Figure 6.**
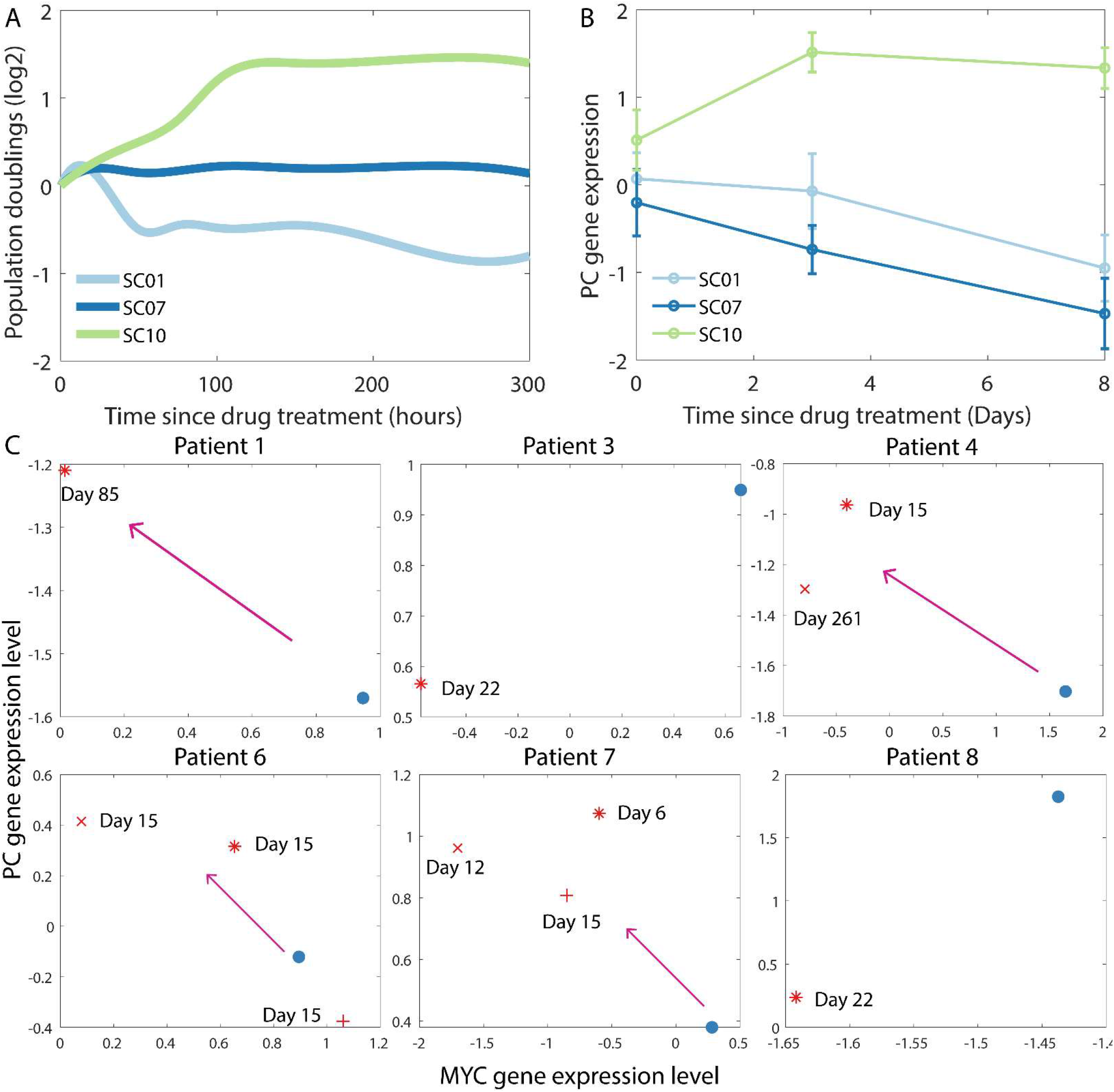
Pyruvate carboxylase (PC) expression levels are altered in melanoma cells in response to BRAF signaling inhibitors. (A) Representative depiction of the response of three different subclones of the SKMEL5 melanoma cell line to treatment with PLX4720, a mutant BRAF inhibitor, as reported by Paudel *et al*. [41] (see Fig. 3 A therein). (B) RNA expression levels of the PC gene (Z-score calculated after log2 normalization) in the three subclones in (A) at 3 days and 8 days post-treatment with PLX4720. The gene expression data was obtained from Jia *et al*. [42]. The error bars indicate the standard deviation. (C) RNA expression levels of MYC and PC genes (Z-score calculated after log2 normalization) in tumor samples obtained from melanoma patients. In each panel, the blue dot indicates the expression levels before treatment. The different red markers in each panel indicate the expression levels in samples obtained at different time points after treatment with BRAF signaling inhibitors. The gene expression data was from downloaded from the Gene Expression Omnibus databse (accession GSE75299). See Song *et al*. [43] for details regarding the patient samples and the BRAF signaling inhibitor administered in each case.

## Discussion

Multiple studies have shown that tumor cells within the primary tumor and those at different stages of metastatic progression exhibit widespread heterogeneity in their metabolic profiles [4,5]. The metabolic phenotype can also co-vary with other functional attributes such as morphology [47], migratory phenotype [32,33], and stemness [48]. Here, using a detailed mechanistic model of some of the key metabolic pathways in tumor cells, we show that variation in ATP usage can be a key driver of the metabolic heterogeneity exhibited by tumor cells. Differences in the availability of nutrients and in the ability of tumor cells to take up these nutrients can further contribute towards the diversity of metabolic profiles. By using a mechanistic modeling approach, our study avoids the difficulties associated with choosing an appropriate objective function and appropriate constraints that plague constraint-based models. Further, our modeling approach captures the effect of feedback loops in metabolic pathways, and we show that these are crucial to understanding the metabolic heterogeneity exhibited by tumor cells. Exploring the possible set of metabolic profiles that can be obtained by varying the reaction velocities in the model instead of fitting the model parameters to a set of experimental observations concerning a specific cell line allowed us to develop a broad framework to analyze diverse tumor metabolic profiles instead of making predictions pertaining only to a given experimental setup. While the framework in the present study does not include all the metabolic pathways active in tumor cells as is usually the case for genome-scale metabolic models [6], the predictions from our framework are far more interpretable and propose key principles that underly the metabolic profiles seen across tumor types. These principles could hold the key to identifying metabolic targets for anti-cancer therapy [5].

Our model predicts that in order to drive large fluxes through anabolic pathways that are crucial to fast proliferation, the cytosolic ATP concentration must be kept below a threshold to maintain the activity of PFK, a key enzyme in the glucose uptake pathway. This can explain why tumor cells, across cancer types, often prefer aerobic glycolysis over oxidative phosphorylation— the ATP yield per glucose molecule from aerobic glycolysis is less than one-tenth of the yield from oxidative phosphorylation [29]. That the inhibition of PFK by high levels of ATP can drive aerobic glycolysis in tumor cells was first postulated by Scholnick *et al*. [49] nearly half a century ago. An important corollary that follows from this prediction is that the preference of tumor cells for aerobic glycolysis versus oxidative phosphorylation can be modulated by the rate of ATP consumption in these cells. We have built upon this corollary to describe how the metabolic state of tumor cells can depend on their migratory phenotype [32,33]. The framework can also be helpful in understanding other behaviors exhibited by tumor and other fast proliferating cells. For example, the PI3K / Akt signaling pathway, which is activated in response to extracellular growth factors in normal, non-cancerous cells [50], transcriptionally upregulates ENTPD5. ENTPD5 promotes proper protein glycosylation and protein folding via a cycle of reactions that convert ATP to ADP [51]. Our model predicts that the resulting increase in ATP consumption will increase the flux through anabolic pathways such as the phospholipids synthesis pathway, facilitating fast proliferation. Knockdown of ENTPD5 has indeed been shown to inhibit cell growth in PTEN-null mouse embryonic fibroblasts [51]. The increased glycolytic flux from upregulated ATP consumption can also account for the observed upregulation of ATPases such as the Na^+^ / K^+^ pump in tumor cells [52–54]. The idea that the operation of the Na^+^ / K^+^ pump can alter the rate of glycolysis was also proposed by Scholnick *et al*. [49]. Note that the model of tumor metabolism described here does not incorporate ATP production from other metabolic pathways including from the *β*-oxidation of fatty acids. Inclusion of *β*-oxidation in the present model could shed light on the interplay between fatty acid metabolism and other cell behaviors including proliferation rates, and further unravel the context-dependence of fast proliferation rates on glycolytic flux.

Recently, Luengo *et al*. [55] have proposed that increased NAD^+^ demand for oxidation as compared to the ATP demand during cell proliferation can drive a preference for aerobic glycolysis in multiple cell lines. In such a scenario, high ATP concentration and low ADP availability, resulting from low ATP usage, inhibits the regeneration of NAD^+^ from the NADH produced during the TCA cycle. The limited NAD^+^ availability limits cell proliferation. In the modeling framework described in the present study, high ATP concentration resulting from low ATP usage also drives a preference for aerobic glycolysis. However, in our framework, the NAD^+^ concentration is not limiting. Rather, it is the anabolic fluxes, such as the flux through the phospholipids synthesis pathway, that become limiting due to the inhibition of PFK by ATP. The focus on increased anabolic fluxes required for cell division allows us to capture the increased glucose uptake and lactate production in tumor cells. Limited NAD^+^ availability cannot account for the increased lactate production since uptake of carbon as glucose and its excretion as lactate is redox neutral overall. The two ideas, one described by Luengo *et al*. [55] and the other proposed in the present study, may be applicable in different scenarios depending on whether NAD^+^ is limiting or not.

Tumor cells, across cancer types, can rely on different growth signaling pathways to drive continued cell proliferation [56,57]. Different growth signaling pathways often activate distinct pathways to drive the same cellular process. For example, in prostate cancer cells, androgen-receptor mediated signaling promotes the uptake of fatty acids from the extracellular environment while PI3K / Akt signaling drives de novo fatty acid synthesis [35]. In the case of breast cancer, receptor-positive breast cancer cells rely primarily on de novo fatty acid synthesis while basal-like, receptor-negative breast cancer cells take up fatty acids from the extracellular environment [58]. The present modeling framework can provide useful insights into the metabolic profiles of cancer cells reliant on different growth signaling pathways. Our model predicts that tumor cells that synthesize fatty acids de novo must increase their glutamine uptake or upregulate the expression of the enzyme PC. While cells taking up glutamine will maintain high lactate production, lactate production will be down-regulated in cells upregulating PC activity. In contrast, tumor cells that can take up fatty acids can rely on these both for ATP generation (via *β*-oxidation) and for anabolic processes. These cells will thus likely have both low glutamine uptake and low glucose uptake. Prostate cancer cells treated with an inhibitor of androgen receptor signaling such as enzalutamide can develop resistance by activating the PI3K / Akt signaling pathway, thereby turning on ***de novo*** fatty acid synthesis [59]. Our modeling framework predicts that the emergence of enzalutamide resistance must be accompanied by a change in the metabolic profile of tumor cells including an increase in glucose uptake. A recent study has shown that the activities of signaling pathways that regulate cell metabolism, including the p38 pathway, are indeed altered by enzalutamide treatment in prostate cancer cells [60].

Multiple studies have shown a tight coupling between cell-type switching and change in the metabolic state of cells, especially in the context of epithelial-mesenchymal plasticity. Bhattacharya *et al*. [61] showed that neural crest cells in a chicken embryo activate aerobic glycolysis at the onset of cell migration and that an increase in glycolytic flux is essential for driving an epithelial to mesenchymal transition. Luo *et al*. [47] have shown that inhibition of glucose uptake via treatment with 2-deoxyglucose leads to an increase in reactive oxygen species (ROS) concentration, causing mesenchymal breast cancer stem cells to switch to a more epithelial state (but not to a fully epithelial state). Our modeling framework suggests that inhibition of glucose uptake will lead to increased reliance on glutamine for ATP production (Fig. 5 A and Fig. 5 B). Apart from playing a key role in anaplerosis and in glucose-independent ATP production, glutamine also drives an anti-oxidant response via the glutathione pathway [62]. Treatment with 2-deoxyglucose could thus decrease the availability of glutamine for the anti-oxidant response. The consequent increase in ROS concentration can upregulate NRF2 which has been shown to stabilize a hybrid epithelial-mesenchymal state [63]. Further, given the low availability of glutamine for the anti-oxidant pathway, epithelial cells would be more susceptible to thioredoxin suppression [47] (thioredoxin mediates a glutamine-independent anti-oxidant response). Incorporating ROS generation and anti-oxidant pathways into the present modeling framework could thus help further our understanding of metabolic dependence in cell-type switching.

Finally, the present model can be extended to develop a population-level framework to analyze the interaction between tumor cells exhibiting different metabolic phenotypes. For example, tumor cells exhibiting glycolysis can cooperate with non-glycolytic tumor cells by excreting lactate which can be taken up by the neighboring cells and metabolized via the TCA cycle [64]. The lactate excreted by glycolytic tumor cells has also been shown to support the metabolic needs of tumor-promoting regulatory T cells, and suppression of the lactate transporter MCT1 in regulatory T cells has been shown to synergize with anti-PD-1 therapy in a mouse model of melanoma [65]. Moreover, tumor cells and immune cells compete for nutrients in the tumor microenvironment and tilting the competition towards tumor-suppressive immune cells can prove to be an effective anti-tumor strategy [38,66]. A population-level extension of the present framework could prove invaluable in developing strategies to modulate the metabolic interplay in the tumor microenvironment to a tumor-suppressive state.

## Materials and Methods

### Cell culture and knockdown of pyruvate carboxylase

MDA-MB-468 breast cancer cells were maintained in DMEM cell culture medium supplemented with 5% FBS, 100 units / ml penicillin, and 100 μg / ml streptomycin. The cell line was authenticated using short tandem repeat profiling by the MD Anderson Cancer Center Cytogenetics and Cell Authentication Core. pGIPZ lentivirus shRNA for pyruvate carboxylase (PC) was purchased from the Cell-Based Assay Screening Service (C-BASS) Core at Baylor College of Medicine. The lentivirus with scrambled (control) or PC-targeting shRNA was infected using the standard protocol [67]. Knockdown efficiency was validated by western blot analysis using specific antibodies.

### Cell Respiratory Assay

Oxygen consumption rate (OCR) was measured using the XFp extracellular flux analyzer (Seahorse Biosciences) as previously described [67] with a minor modification. Cell Mito Stress kit (Seahorse Biosciences) was used for the assay and basal OCR was calculated by Report Generator Version 4.03 (Seahorse Biosciences).

### High-resolution Nuclear Magnetic Resonance (NMR) Spectroscopy

Metabolites were extracted from cell pellets using 3 mL 2:1 methanol-water solvent and 0.5 ml of lysing beads (Lysing Matrix D from MP Biomedicals, LLC). The mixture was vortexed and freeze-thawed to extract the metabolites. This was followed by centrifugation at 4,000 RPM for 10 minutes to remove debris, and rotary evaporation and overnight lyophilization to remove the solvent. The samples were prepared for NMR spectroscopy by dissolving the lyophilized sample in 800 μL of ^2^H_2_O following the centrifugation at 10,000 RPM for 5 minutes to remove any debris that remained. Finally, 600 μL of the sample, with the addition of 40 μL of 8 mM NMR reference compound 2, 2-dimethyl-2-silapentane-5-sulfonate (DSS), was used for NMR spectroscopy. Final concentration of DSS in the sample was 0.5 mM. The standard one dimensional 1H with water suppression during relaxation time was used to acquire the data. All 1D spectra were acquired with 32K time domain points, 2 seconds acquisition time, 256 transients, 64 receiver gain, 16 ppm spectral width, and 6 second relation delay on a Bruker NMR spectrometer operating at 500 MHz proton resonance frequency equipped with cryogenically cooled triple resonance (^1^H, ^13^C, ^15^N) TXI probe [68]. The data was processed using Topspin 3.1 and resonances were identified using Chenomx NMR suite 7.0 from Chenomx Inc and the human metabolic database (HMDB) [69].

## Author Contributions

Conceptualization, S.T. and H.L.; Methodology, S.T. and J.H.K.; Investigation, S.T, J.H.K, S.P, and P.K.B.; Writing-Original Draft Preparation, S.T.; Writing-Review & Editing, S.T., J.H.K, B.A.K, and H.L.; Supervision, B.A.K. and H.L.

## Supporting Information

**S1 Text:** Includes a detailed description of the mathematical model of tumor metabolism presented in the study, along with the assumptions underlying model construction

