## Supplementary Text for "A Mechanistic Modeling Framework Reveals the Key Principles Underlying Tumor Metabolism"

### A mathematical model of tumor metabolism

We used ordinary differential equations to model the dynamical behavior of the system of metabolic reactions. The different metabolites in our model and the corresponding abbreviated metabolite names are listed in Table S1. The mathematical expressions describing the activities of the different enzymes are available as MATLAB code which can be downloaded from https://github.com/st35/cancer-metabolism. The MATLAB code also includes the values of the kinetic parameters for each enzyme. The code for generating the different figures included in the manuscript is also available at the same link.

The rate of change of concentration of a metabolite is given by the rates of production and consumption of the metabolite. Below, we list the differential equations governing the concentrations of the different metabolites. The terms on the right hand sides of these equations represent the fluxes through the different enzymes as listed in Table S2. Metabolites whose concentrations were kept fixed and the corresponding concentrations are listed in Table S3.

$$\frac{d\left[C_{GLC}\right]}{dt} = r_{GLUT} - r_{HK} \qquad \text{Eq. 1}$$

$$\frac{d\left[C_{G6P}\right]}{dt} = r_{HK} - r_{GPI} - r_{G6PD} \qquad \text{Eq. 2}$$

$$\frac{d\left[C_{F6P}\right]}{dt} = r_{GPI} - r_{PFK} \qquad \text{Eq. 3}$$

$$\frac{d\left[C_{F16BP}\right]}{dt} = r_{PFK} - r_{ALD} \qquad \text{Eq. 4}$$

$$\frac{d\left[C_{DHAP}\right]}{dt} = r_{ALD} - r_{TPI} - r_{G3PD} \qquad \text{Eq. 5}$$

$$\frac{d\left[C_{GAP}\right]}{dt} = r_{ALD} + r_{TPI} - r_{GAPDH} \qquad \text{Eq. 6}$$

$$\frac{d\left[C_{13BPG}\right]}{dt} = r_{GAPDH} - r_{PGK} \qquad \text{Eq. 7}$$

$$\frac{d\left[C_{3PG}\right]}{dt} = r_{PGK} - r_{PGM} \quad \text{Eq. 8}$$

$$\frac{d\left[C_{2PG}\right]}{dt} = r_{PGM} - r_{ENO} \quad \text{Eq. 9}$$

$$\frac{d\left[C_{PEP}\right]}{dt} = r_{ENO} - r_{PK} \quad \text{Eq. 10}$$

$$\frac{d\left[C_{PYR}\right]}{dt} = r_{PK} - r_{LDH} - r_{PIM} \quad \text{Eq. 11}$$

$$\frac{d\left[M_{PYR}\right]}{dt} = r_{PIM} - r_{PDH} \quad \text{Eq. 12}$$

$$\frac{d\left[M_{ACCOA}\right]}{dt} = r_{PDH} - r_{CIT} \quad \text{Eq. 13}$$

$$\frac{d\left[M_{CIT}\right]}{dt} = r_{CIT} - r_{ACO} \quad \text{Eq. 14}$$

$$\frac{d\left[M_{ICIT}\right]}{dt} = r_{ACO} - r_{IDH} \quad \text{Eq. 15}$$

$$\frac{d\left[M_{AKG}\right]}{dt} = r_{IDH} - r_{AKGD} \quad \text{Eq. 16}$$

$$\frac{d\left[M_{SCOA}\right]}{dt} = r_{AKGD} - r_{SCOAS} \quad \text{Eq. 17}$$

$$\frac{d\left[M_{SUC}\right]}{dt} = r_{SCOAS} - r_{SDH} \quad \text{Eq. 18}$$

$$\frac{d\left[M_{FUM}\right]}{dt} = r_{SDH} - r_{FUM} \quad \text{Eq. 19}$$

$$\frac{d\left[M_{MAL}\right]}{dt} = r_{FUM} - r_{MDH} \quad \text{Eq. 20}$$

$$\frac{d\left[M_{OAA}\right]}{dt} = r_{MDH} - r_{CIT} \quad \text{Eq. 21}$$

$$\frac{d\left[M_{NADH}\right]}{dt} = r_{PDH} + r_{IDH} + r_{AKGD} + r_{MDH} - r_{Resp} \quad \text{Eq. 22}$$

$$\frac{d\left[M_{\Delta\psi}\right]}{dt} = 10r_{Resp} - 3r_{ATP} - r_{ANT} - r_{Leak} \quad \text{Eq. 23}$$

$$\frac{d\left[M_{ATP}\right]}{dt} = r_{ATP} - r_{ANT} \quad \text{Eq. 24}$$

$$\frac{d\left[M_{ADP}\right]}{dt} = r_{ANT} - r_{ATP} \quad \text{Eq. 25}$$

$$\frac{d[C_{ATP}]}{dt} = r_{ANT} - r_{ATP\,Use} + r_{PGK} + r_{PK} - r_{HK} - r_{PFK} \quad \text{Eq. 26}$$

$$\frac{d[C_Q]}{dt} = r_{SLC1A5} - r_{SLC1A5\_var} \quad \text{Eq. 27}$$

$$\frac{d[M_Q]}{dt} = r_{SLC1A5\_var} - r_{GLS} \quad \text{Eq. 28}$$

$$\frac{d[M_{GLU}]}{dt} = r_{GLS} - r_{GLUD} \quad \text{Eq. 29}$$

$$\frac{d[C_{CIT}]}{dt} = r_{SLC25A1} - r_{ACLY} \quad \text{Eq. 30}$$

$$\frac{d[C_{ACCOA}]}{dt} = r_{ACLY} - r_{ACC} \quad \text{Eq. 31}$$

### Fig. 1 B-D

For the analysis shown in Fig. 1 B-D, we assumed that any ATP produced via glycolysis or the tricarboxylic acid (TCA) cycle is consumed by cellular processes, keeping the concentration of ATP in the cytoplasm and the mitochondria fixed. To obtain the results shown in these figures, we integrated Eq. 1-21. The $NAD^+$:NADH ratio in both the mitochondria and the cytoplasm was also kept fixed. The mathematical expressions describing the activities of the different enzymes and the relevant kinetic parameters were obtained from the study by Mulukutla *et al.* [1] and from the study by Wu *et al.* [2].

### Fig. 1 F, Fig. 1 G, and Fig. 2

Reactions involved in glycolysis and the TCA cycle were modeled using Eq. 1-21. The concentrations of ATP in the mitochondria and in the cytoplasm were no longer kept fixed, and were governed by Eq. 24 and Eq. 26, respectively. ATP production from oxidative phosphorylation and the electron transport chain was modeled using the coarse-grained approached described previously by Nazaret *et al.* [3] (Eq. 22-26). Briefly, the NADH produced from the TCA cycle reactions is used to maintain a transmembrane potential ($M_{\Delta\psi}$ in Eq. 23) by pumping protons from the inner mitochondrial matrix to the outside (represented by the term $r_{Resp}$ in Eq. 22 and Eq. 23). The oxidation of one NADH molecule to $NAD^+$ provides the energy for pumping 10 protons from the inner mitochondrial matrix to the outside, across the inner mitochondrial membrane (consequently, the factor of 10 in the first term of Eq. 23). This is an oxygen-dependent process and we modeled the dependence of the rate of NADH oxidation on the oxygen availability by modifying the mathematical expression in Nazaret *et al.* [3] (see equation 5 therein):

$$r_{Resp} = v_{resp}\left(\frac{M_{O_2}}{K_{O_2} + M_{O_2}}\right)$$

where $v_{resp}$ is the expression for the rate of NADH oxidation proposed by Nazaret *et al.* [3], $M_{O_2}$ is the concentration of oxygen, and $K_{O_2}$ is a constant. The transmembrane potential across the inner mitochondrial membrane is utilized by the ATP synthase to convert ADP to ATP. For synthesizing one ATP molecule, the energy supplied by three protons entering the inner mitochondrial space from the cytosol is needed (hence a factor of 3 in the second term of Eq. 23). The term $r_{ANT}$ captures the exchange of one ATP from the inner mitochondrial space for one ADP from the outside via the adenine nucleotide translocator. Finally, $r_{Leak}$ describes the leakage of protons across the inner mitochondrial membrane, contributing to the loss of the transmembrane potential over time. The mathematical expressions for the different right hand side terms in Eq. 22-26 can be obtained from the MATLAB code.

The rate of ATP use by cells, represented by the term $r_{ATP\ Use}$ in Eq. 26, increases with the availability of ATP and decreases at high ADP concentrations:

$$r_{ATP\ Use} = \frac{V_{ATP\ Use}\left(\frac{C_{ATP}}{k_{ATP\ Use}}\right)}{1 + \left(\frac{C_{ATP}}{k_{ATP\ Use}}\right) + \left(\frac{C_{ADP}}{k_{ADP}}\right)}$$

Here, $C_{ATP}$ and $C_{ADP}$ are the cytoplasmic ATP and ADP concentrations, respectively. Varying ATP requirements in tumor cells were modeled by multiplying $V_{ATP\ Use}$ with a factor ($V_{ATP\ Use} = F \times V^0_{ATP\ Use}$). It is this multiplicative factor $F$ that is shown along the horizontal axes in the panels of Fig. 2. In Fig. 1 F and Fig. 1 G, $F = 0.1$.

Note that the total concentration of ADP and ATP is kept constant, both in the cytoplasm ($C_{ADP} + C_{ATP} =$ constant) and in the mitochondria ($M_{ADP} + M_{ATP} =$ constant).

### Synthesis of fatty acids

In our model, fatty acids are synthesized in the cytoplasm from the citrate exported out of the mitochondria by the mitochondrial citrate carrier (SCL25A1). During the NADPH-dependent reductive carboxylation of $\alpha$-ketoglutarate, $\alpha$-ketoglutarate is directly converted to citrate. To incorporate these processes into the model, Eq. 14 was modified:

$$\frac{d[M_{CIT}]}{dt} = r_{CIT} - r_{ACO} - r_{SCL25A1} + r_{IDH\_Rev}$$

Here, $r_{SLC25A1}$ represents the export of citrate from the mitochondria and $r_{IDH\_Rev}$ represents the flux through the reductive carboxylation reaction. Our model includes only the first two steps in the fatty acid synthesis pathway— the conversion of citrate to acetyl-CoA by the enzyme ACLY and the conversion of acetyl-CoA to malonyl-CoA by the enzyme ACC (see Eq. 30 and Eq. 31). Note that our model does not incorporate the production for acetyl-CoA in the cytoplasm from other sources such as acetate.

### Glutaminolysis

First, since the conversion of glutamate to $\alpha$-ketoglutarate generates one molecule of NADH (from $NAD^+$), Eq. 22 was modified as shown below:

$$\frac{d[M_{NADH}]}{dt} = r_{PDH} + r_{IDH} + r_{AKGD} + r_{MDH} + r_{GLUD} - r_{Resp}$$

Here, $r_{GLUD}$ represents the flux through the enzyme glutamate dehydrogenase.

Next, for the system of metabolic reactions in our model to exhibit a bounded steady state, the rate of production of $\alpha$-ketoglutarate from glutamate must exactly equal the flux through the fatty acid synthesis pathway. This is because the role of glutamine in anaplerosis is to simply replenish the carbon being siphoned off from the TCA cycle as citrate for fatty acid synthesis. To ensure that a bounded steady state exists, we added an $\alpha$-ketoglutarate transport term to Eq. 16. This ensures that any excess $\alpha$-ketoglutarate generated from glutamate is transported from the mitochondrial matrix to the cytosol ($\alpha$-ketoglutarate may be used for biosynthetic processes in the cytoplasm and as a cofactor for histone modifiers in the nucleus [4]). Thus, Eq. 16 becomes

$$\frac{d[M_{AKG}]}{dt} = r_{IDH} - r_{AKGD} + r_{GLUD} - r_{IDH\_Rev} - r_{SLC25A11}$$

Here, $r_{GLUD}$ represents the conversion of glutamate to $\alpha$-ketoglutarate, $r_{IDH\_Rev}$ represents the flux through the reductive carboxylation reaction, and $r_{SLC25A11}$ represents the export of $\alpha$-ketoglutarate from the mitochondria to the cytoplasm (via the mitochondrial $\alpha$-ketoglutarate carrier SLC25A11).

While constructing genome-scale metabolic models of tumor cells, reactions in addition to those involving tumor-specific enzymes must be included in order to ensure that the resultant metabolic model is consistent and obeys stoichiometric and mass balance constraints, among others (for example, see Folger *et al.* [5]). Our inclusion of the $\alpha$-ketoglutarate transport term in Eq. 16 to ensure that the metabolic model has a bounded state is motivated by the previous work on genome-scale metabolic models. However, since the focus of this study is on keeping the model size small, we do not incorporate all the additional metabolic pathways that may be needed for the metabolite concentrations in the system to be bounded at steady state when incorporating glutaminolysis reactions.

### Pyruvate carboxylase-mediated anaplerosis

Pyruvate carboxylase-mediated anaplerosis involves the conversion of pyruvate to oxaloacetate in the mitochondria. The process consumes one molecule to ATP per pyruvate molecule converted. To incorporate this reaction into our model, we modified Eq. 12, Eq. 21, and Eq. 24:

$$\frac{d\left[M_{PYR}\right]}{dt} = r_{PIM} - r_{PDH} - r_{PC}$$

$$\frac{d\left[M_{OAA}\right]}{dt} = r_{MDH} - r_{CIT} + r_{PC}$$

$$\frac{d\left[M_{ATP}\right]}{dt} = r_{ATP} - r_{ANT} - r_{PC}$$

Here, $r_{PC}$ represents the flux through the enzyme pyruvate carboxylase (PC).

### Malic enzyme 2-mediated anaplerosis

Malic enzyme 2-mediated anaplerosis involves the conversion of malate to pyruvate in the mitochondria, a process accompanied by the conversion of one $NAD^+$ molecule to NADH. To incorporate this reaction into our model, we modified Eq. 12, Eq. 20, and Eq. 22:

$$\frac{d\left[M_{PYR}\right]}{dt} = r_{PIM} - r_{PDH} - r_{PC} + r_{ME2}$$

$$\frac{d\left[M_{MAL}\right]}{dt} = r_{FUM} - r_{MDH} - r_{ME2}$$

$$\frac{d\left[M_{NADH}\right]}{dt} = r_{PDH} + r_{IDH} + r_{AKGD} + r_{MDH} + r_{GLUD} + r_{ME2} - r_{Resp}$$

Here, $r_{ME2}$ represents the flux through malic enzyme 2 (ME2).

### Model assumptions

In the present study, our focus is on analyzing a mechanistic model of some of the key metabolic pathways active in tumor cells. Given the difficulty of determining the mathematical expressions describing the kinetic behavior of different enzymes and the relevant kinetic parameters, we focused on a relatively small set of key reactions. As compared to genome-scale metabolic models of tumor metabolism studied previously (reviewed in [6]) which can include thousands of reactions, the number of chemical reactions in our model is miniscule. Consequently, our model involves multiple assumptions concerning how the concentrations of some of the metabolites are dealt with, and regarding some metabolic reactions.

First, we assume that the $NAD^+$:NADH ratio in the cytoplasm remains constant. Conversion of one glucose molecule to two pyruvate molecules converts two molecules of $NAD^+$ to two molecules of NADH. If the pyruvate is then converted to lactate, NADH is converted back to $NAD^+$. Thus, lactate production can maintain a constant $NAD^+$:NADH ratio in the cytoplasm. In the case where pyruvate enters the TCA cycle, the cytoplasmic NADH produced during the conversion of glucose to pyruvate can be converted back to $NAD^+$ via the glycerol-3-phosphate shuttle [7]. While this will once again maintain a constant cytoplasmic $NAD^+$:NADH ratio as we have assumed, our model does not include the reactions of the glycerol-3-phosphate shuttle and does not account for the ATP produced by this process (3 or 5 ATPs per molecule of glucose [8]). Our model also does not account for $FADH_2$ production in the mitochondria via the TCA cycle. Thus, overall, our model underestimates, to some extent, the ATP production from the TCA cycle and oxidative phosphorylation. Developing the model further to exactly account for ATP production from these processes will however not qualitatively change any of the model predictions— the ATP production per glucose molecule when glucose is used to drive the TCA cycle remains more than 10 fold higher than when glucose is used for lactate production in our model.

Second, the concentration of malate in the cytoplasm was kept fixed. This was despite the fact that our model includes the transport of citrate from the inner mitochondrial matrix to the cytoplasm, and this transport process is coupled with the transport of malate from the cytoplasm into the mitochondria. As evident from Eq. 20, the malate concentration in the mitochondria in our model is also unaffected by the export of citrate from the mitochondria to the cytoplasm. Here, we consider the operation of a cascade of mitochondrial SLC25 transporters: SLC25A1 which exports citrate in exchange for cytoplasmic malate, followed by SLC25A10 which exports malate in exchange for cytoplasmic phosphate [9]. Thus, the overall reaction in the mitochondria is the export of citrate and import of phosphate, with the malate concentration in the inner mitochondrial matrix remaining unaffected by citrate export. Therefore, we did not include a term representing citrate export in Eq. 20. Note that when citrate is converted to acetyl-CoA in the cytoplasm by the enzyme ACLY, malate is generated as a byproduct. Thus, overall, for each citrate molecule exported from the mitochondrial matrix to the cytoplasm, one molecule of malate is added to the cytoplasm. We assume that the malate generated in the cytoplasm by the conversion of citrate to acetyl-CoA is used up in other metabolic pathways, maintaining a constant cytoplasmic malate concentration.

Third, our model incorporates only the first reaction involved in the two key anabolic processes— the ribose synthesis pathway and the phospholipids synthesis pathway. For the fatty acid synthesis pathway, only the first two reactions are included. The assumption here is that the downstream reactions in these pathways are not limiting and that there is no feedback from downstream metabolites to the first reactions in these pathways.

Fourth, as mentioned earlier, to ensure that the system of metabolic reactions exhibits a bounded steady state, we assume that any excess $\alpha$-ketoglutarate generated from glutamate (excess $\alpha$-ketoglutarate left after the carbon siphoned off from the TCA cycle as citrate has been replenished) is exported out of the mitochondria. This is a simplifying assumption which relies on the existence of a feedback mechanism which couples the rate of $\alpha$-ketoglutarate generation from glutamate to the rate at which carbon is being removed from the TCA cycle for biosynthetic processes. Such a feedback mechanism likely involves one of the metabolic processes that is not included in our present modeling framework.

### Comment on model construction

A key challenge in constructing mechanistic models of metabolic pathways is that the data concerning enzyme kinetics available in the literature are often not compatible [2]— the expressions describing enzyme kinetics as well as the relevant kinetic parameters used in the present study were obtained from multiple previous studies which themselves relied on results from experiments conducted in different tissue types, in cells from different species, and under a varied set of conditions [1,2,10–12]. This is in addition to the unavailability of the relevant data for some reactions. Thus, one must be careful while making quantitative predictions from such models regarding the concentrations of different metabolites in cells. Mechanistic modeling of metabolism presents an additional challenge. The mathematical expressions describing the activities of the different enzymes and the relevant kinetic parameters must be such that all metabolic fluxes balance. Otherwise, the system will not exhibit a bounded, non-trivial steady state. The metabolic fluxes may not balance when the mathematical expressions and kinetic parameters taken from diverse sources are put together into a single mathematical model. This is possible (a) when the available data are from studies conducted in different cell types, in different species, and under different conditions (which is usually the case as mentioned above), and (b) if the model is missing certain key metabolic reactions that will allow for a bounded steady state for a given parameter set.

In the present study, our goal was not to make quantitative predictions regarding the concentrations of the different metabolites or to fit the model behavior to data from any specific experimental setup. This allowed us to focus on analyzing the behavior of a few key metabolic pathways in tumor cells instead of working towards a more "complete" model as may be essential for making precise quantitative predictions. To ensure that the system consisting of a limited set of metabolites and reactions in our modeling framework exhibits a bounded steady state, we manually adjusted the values of some of the kinetic parameters obtained from previous studies. All the kinetic parameters and the MATLAB code used to generate the figures in the manuscript are available online (https://github.com/st35/cancer-metabolism). It is possible, and very likely, that there are numerous other sets of adjusted kinetic parameters that will allow for the existence of a bounded steady state in our model. The qualitative predictions we make here using a specific set of kinetic parameters will not change for a different choice of parameters provided a key condition holds— the cytoplasmic ATP concentration becomes high enough to inhibit PFK activity before the ADP concentration becomes too low for the continued operation of oxidative phosphorylation and the electron transport chain (note that the sum of ATP and ADP concentrations remains constant). Even when this condition is not met, low ATP consumption would still be a driver of a preference for the Warburg effect but instead of high ATP concentration inhibiting glucose uptake at the level of PFK, it will be low ADP concentration inhibiting glucose oxidation via the TCA cycle (and thus promoting lactate production). Behavior similar to this has recently been reported by Luengo *et al.* [13].

**Table S1** The metabolites in our model, abbreviated metabolite names used in the equations, and the variable names used in the MATLAB code.

| Metabolite name | Corresponding term in Eq. 1-31 | Corresponding variable name in MATLAB code |
|---|---|---|
| Cyto. glucose | $C_{GLC}$ | C_GLC |
| Cyto. glucose-6-phosphate | $C_{G6P}$ | C_G6P |
| Cyto. fructose-6-phosphate | $C_{F6P}$ | C_F6P |
| Cyto. fructose-1-6-biphosphate | $C_{F16BP}$ | C_F16BP |
| Cyto. dihydroxyacetone phosphate | $C_{DHAP}$ | C_DHAP |
| Cyto. glyceraldehyde-3-phosphate | $C_{GAP}$ | C_GAP |
| Cyto. 1, 3-biphosphoglycerate | $C_{13BPG}$ | C_13BPG |
| Cyto. 3-phosphoglycerate | $C_{3PG}$ | C_3PG |
| Cyto. 2-phosphoglycerate | $C_{2PG}$ | C_2PG |
| Cyto. phosphoenolpyruvate | $C_{PEP}$ | C_PEP |
| Cyto. pyruvate | $C_{PYR}$ | C_PYR |
| Mito. pyruvate | $M_{PYR}$ | M_PYR |
| Mito. acetyl-CoA | $M_{ACCOA}$ | M_ACCOA |
| Mito. citrate | $M_{CIT}$ | M_CIT |
| Mito. isocitrate | $M_{ICIT}$ | M_ICIT |
| Mito. $\alpha$-ketoglutarate | $M_{AKG}$ | M_AKG |
| Mito. succinyl-CoA | $M_{SCOA}$ | M_SCOA |
| Mito. succinate | $M_{SUC}$ | M_SUC |
| Mito. fumarate | $M_{FUM}$ | M_FUM |
| Mito. malate | $M_{MAL}$ | M_MAL |
| Mito. oxaloacetate | $M_{OAA}$ | M_OAA |
| Mito. NADH | $M_{NADH}$ | M_NADH |
| Mito. ATP | $M_{ATP}$ | M_ATP |
| Mito. ADP | $M_{ADP}$ | M_ADP |
| Cyto. ATP | $C_{ATP}$ | C_ATP |
| Cyto. glutamine | $C_{Q}$ | C_Q |
| Mito. glutamine | $M_{Q}$ | M_Q |
| Mito. glutamine | $M_{GLU}$ | M_GLU |
| Cyto. citrate | $C_{CIT}$ | C_CIT |
| Cyto. acetyl-CoA | $C_{ACCOA}$ | C_ACCOA |

**Table S2** The enzymes in our model, terms representing the fluxes through the enzymes in the equations, and the variable names representing the fluxes in the MATLAB code.

| Enzyme name | Corresponding term in Eq. 1-31 | Corresponding variable name in MATLAB code |
|---|---|---|
| Glucose transporter | $r_{GLUT}$ | r_GLUT |
| Hexokinase | $r_{HK}$ | r_HK |
| Glucose phosphate isomerase | $r_{GPI}$ | r_GPI |
| Glucose-6-phosphate dehydrogenase | $r_{G6PD}$ | r_G6PD |
| Phosphofructokinase | $r_{PFK}$ | r_PFK |
| Aldolase | $r_{ALD}$ | r_ALD |
| Triose phosphate isomerase | $r_{TPI}$ | r_TPI |
| Glycerol-3-phosphate dehydrogenase | $r_{G3PD}$ | r_G3PD |
| Glyceraldehyde-3-phosphate dehydrogenase | $r_{GAPDH}$ | r_GAPDH |
| Phosphoglycerate kinase | $r_{PGK}$ | r_PGK |
| Phosphoglycerate mutase | $r_{PGM}$ | r_PGM |
| Enolase | $r_{ENO}$ | r_ENO |
| Pyruvate kinase | $r_{PK}$ | r_PK |
| Lactate dehydrogenase | $r_{LDH}$ | r_LDH |
| Mitochondrial pyruvate transporter | $r_{PIM}$ | r_PIM |
| Pyruvate dehydrogenase | $r_{PDH}$ | r_PDH |
| Citrate synthase | $r_{CIT}$ | r_CIT |
| Aconitase | $r_{ACO}$ | r_ACO |
| Isocitrate dehydrogenase | $r_{IDH}$ | r_IDH |
| $\alpha$-ketoglutarate dehydrogenase | $r_{AKGD}$ | r_AKGD |
| Succinyl-CoA synthetase | $r_{SCOAS}$ | r_SCOAS |
| Succinate dehydrogenase | $r_{SDH}$ | r_SDH |
| Fumarase | $r_{FUM}$ | r_FUM |
| Malate dehydrogenase | $r_{MDH}$ | r_MDH |
| Glutamine transporter (cell membrane) | $r_{SLC1A5}$ | r_SLC1A5 |
| Mitochondrial glutamate transporter | $r_{SLC1A5\_var}$ | r_SLC1A5_var |
| Glutaminase | $r_{GLS}$ | r_GLS |
| Glutamate dehydrogenase | $r_{GLUD}$ | r_GLUD |
| Mitochondrial citrate-malate antiport | $r_{SLC25A1}$ | r_SLC25A1 |
| Acetyl-CoA lyase | $r_{ACLY}$ | r_ACLY |

| | | |
|---|---|---|
| Acetyl-CoA carboxylase | $r_{ACC}$ | r_ACC |

**Table S3** Metabolites with fixed concentrations in our modeling framework.

| **Metabolite** | **Corresponding variable name in MATLAB code** | **Fixed concentration (mM)** |
|---|---|---|
| Cyto. 2, 3-biphosphoglycerate | C_23BPG | 3.0 |
| Cyto. glutathione | C_GSH | 2.57 |
| Cyto. $Mg^{+}$ | C_Mg | 0.7 |
| Cyto. AMP | C_AMP | 0.03 |
| Cyto. phosphate | C_Pi | 2.5 |
| Cyto. $NAD^{+}$ | C_NAD | 0.299 |
| Cyto. NADH | C_NADH | 0.001 |
| Cyto. pH | C_pH | 7.3 |
| Cyto. alanine | C_ALA | 0.2 |
| Cyto. NADP | C_NADP | 0.0014 |
| Cyto. NADPH | C_NADPH | 0.0643 |
| Mito. coenzyme A | M_COASH | 0.099 |
| Mito. carbon dioxide | M_CO2 | 21.4 |
| Cyto. coenzyme A | C_COA | 0.12 |
| Mito. phosphate | M_Pi | 2.44 |
| Cyto. oxaloacetate | C_OAA | 1.0 |
| Mito. GDP | M_GDP | 0.08 |
| Mito. GTP | M_GTP | $8 \times 10^{-4}$ |
| Mito. ubiquinol | M_COQ | 4.8 |
| Mito. ubiquinone | M_COQH2 | $2.45 \times 10^{-4}$ |
| Mito. ammonia | M_NH3 | $2.4 \times 10^{-3}$ |
| Cyto. ADP + ATP | C_ATP_Tot | 4.0 |
| Mito. $NAD^{+}$ + NADH | M_NAD_Tot | 0.3 |
| Mito. NADP | M_NADP | 0.0014 |
| Mito. NADPH | M_NADPH | 0.0643 |
